# *Pseudomonas aeruginosa* mediates PqsA-dependent iron regulation of the RsmY and RsmZ sRNAs in static conditions

**DOI:** 10.1101/2022.06.23.497436

**Authors:** Rhishita Chourashi, Amanda G. Oglesby

## Abstract

*Pseudomonas aeruginosa* is an opportunistic Gram-negative pathogen that causes acute and chronic lung infection in compromised hosts. During infection, the host innate immune system restricts iron to limit microbial growth. In response, *P. aeruginosa* induces expression of numerous virulence genes. Recently, our lab showed that some virulence factors are responsive to iron limitation in static but not shaking growth conditions, the former of which is likely to mimic growth in the chronically-infected lung. One of these novel iron-responsive factors is the HSI-2-type six secretion system (T6SS), which is also induced during chronic infection. Iron regulation of T6SS was partially impacted by deletion of the iron-responsive PrrF sRNA and completely dependent upon the Pseudomonas quinolone signal (PQS) biosynthetic gene *pqsA*. Here, we analyzed the impact of iron on the expression of two small regulatory RNAs (sRNAs), RsmY and RsmZ, that activate expression of T6SS by sequestering the RsmA translation inhibitor. Our results demonstrate that iron starvation induces expression of RsmY and RsmZ in static but not shaking cultures. We further show that this induction occurs through the *rsmY* and *rsmZ* promoters and is dependent upon PqsA. We identified interrupted palindromes in the *rsmY* and *rsmZ* promoters as putative PqsR binding sites, and disruption of these sites eliminated iron-dependent regulation of *rsmY* and *rsmZ* promoter activity. To determine if iron-dependent regulation of the Rsm sRNAs is likely responsible for iron regulation of HSI-2 T6SS, we constructed translational and transcriptional reporters of the *hsiA2* T6SS gene. Analysis of these reporters revealed robust PqsA-mediated iron regulation of the transcriptional reporter, as well as modest PrrF-dependent iron regulation of the translational reporter. Taken together, our results show novel iron regulatory pathways that are promoted by static growth, highlighting the importance of studying regulatory mechanisms in static communities that are likely more representative of chronic *P. aeruginosa* infections.

**IMPORTANCE:** Iron is a central component of various bacterial metabolic pathways making it an important host acquired nutrient for pathogens to establish infection. Previous iron regulatory studies primaried relied on shaking bacterial cultures; while these ensure cultural homogeneity they do not reflect growth conditions during infection. We recently showed that static growth of *Pseudomonas aeruginosa* promotes iron-dependent regulation of a type six secretion system (T6SS), a virulence factor that is induced during chronic infections. In the current study, we found that static growth also promotes iron-dependent regulation of the RsmY and RsmZ sRNAs, which are global regulators that affect T6SS during chronic *P. aeruginosa* lung infection. Hence, our work demonstrates the Rsm sRNAs as potential effectors of iron regulation during static growth that may also be relevant in chronic infection.

## INTRODUCTION

*Pseudomonas aeruginosa* is a ubiquitous Gram-negative pathogen that causes acute to chronic opportunistic lung infections, due in large part to its diverse metabolic potential and ability to quickly adapt to surrounding conditions and resources (1, 2). *P. aeruginosa* commonly colonizes the lungs of individuals with the hereditary condition cystic fibrosis (CF) in the midst of a diverse microbiota, and eventually shifts to chronic carriage that can persist for decades (3-6). Over time *P. aeruginosa* becomes the predominant bacteria in CF lungs and adapts to the changing environment through several mechanisms that include: i) conversion to a mucoid phenotype; ii) shift from dependency on ferric iron acquisition via siderophores to ferrous and heme acquisition; iii) changes in quorum sensing, with mutations accumulating in the genes for the LasR system while the RhlR and PQS pathways become more active; iv) downregulation of the type three secretion system (T3SS) coupled with increased expression of type six secretion systems (T6SS) (4, 7-16). These adaptations are driven by changes in virulence regulatory pathways that are key to *P. aeruginosa*’s ability to establish chronic infection.

The shift from acute to chronic *P. aeruginosa* virulence in the CF lung is primarily regulated by the GacS/GacA two component system (TCS). GacA, the response regulator of this system, positively regulates production of the RsmY and RsmZ small regulatory RNAs (sRNAs), which themselves control the expression of various virulence genes (17-20). The Rsm sRNAs affect gene expression by sequestering the RsmA RNA-binding protein, which binds to and alters translation efficiency of mRNAs (20). RsmA binds on or near the Shine-Dalgarno sequences of mRNAs encoding three distinct T6SSs (HSI-1, HSI-2 and HSI-3), thus blocking their translation. Thus RsmY and RsmZ prevent RsmA from blocking translation of T6SS mRNAs (20). GacS activation of GacA is inhibited by a distinct sensor kinase, RetS, the gene for undergoes mutational inactivation in many CF isolates of *P. aeruginosa* (21). RetS inactivation indirectly facilitates activation of T6SS during chronic infection by allowing GacSA activation of the Rsm sRNAs. T6SS imparts contact-dependent interactions with bacterial and eukaryotic cells to facilitate fitness of *P. aeruginosa* in the diverse CF lung environment (22). The expression of T3SS genes are inversely down-regulated via the RetS-GacSA-RsmYZ regulatory pathway during chronic infection, underlying the role of T3SS in acute infections (17). Therefore, RetS-GacSA-RsmYZ functions as a central switch from acute (typified by T3SS) to chronic (typified by T6SS) infection phenotypes.

We recently showed that expression of HSI-2 T6SS is also induced upon iron starvation (23). Iron is critical for *P. aeruginosa* survival and virulence, and multiple studies have established iron regulatory effects on virulence gene expression (24). For example, iron starvation in the host induces expression of the PrrF sRNAs (25), which in turn can promote the production of a large family of 2-alkyl-4-quinolone metabolites (AQs), including the Pseudomonas quinolone signal (PQS) (26). The production of PQS is dependent on the *pqsABCDE* operon, the products of which use anthranilate as a precursor for AQ biosynthesis (27). The PrrF sRNAs promote AQ production by inhibiting the translation of the mRNA encoding the AntABC anthranilate degrading enzymes (26, 27). This results in the sparing of anthranilate that can then be used as a precursor for AQ synthesis. PQS is responsible for regulating genes for the production of pyocyanin and elastase, which are virulence factors of *P. aeruginosa*, as well as T6SS (28-31). Our lab previously established that induction of the HSI-2 T6SS by iron starvation is dependent on PqsA and partially dependent on PrrF (23). Interestingly, this regulation was only observed when *P. aeruginosa* strain PAO1 was grown in static conditions, yet the underlying mechanism for static-dependent iron regulation of T6SS remains unclear.

In the current study, we examine whether regulatory factors in the Gac-Rsm pathway are involved in iron regulation of T6SS. We show that iron starvation induces expression of the RsmY and RsmZ sRNAs in static but not shaking conditions. We further show that this regulation is i) dependent upon the *rsmY* and *rsmZ* promoters, ii) completely dependent upon *pqsA*, and iii) partly dependent on the *prrF* locus. These findings led us to determine whether iron regulation of HSI-2 T6SS genes occurred transcriptionally or post-transcriptionally, the latter of which would be consistent with a role for the Rsm sRNAs. Interestingly, our data indicate that iron, PrrF, and AQ-dependent regulation of the T6SS gene *hsi-2* occurs in a promoter-dependent manner. While we have yet to observe evidence for post-transcriptional iron-dependent regulation of T6SS, a thorough examination of how iron affects individual genes and proteins in the HSI-2 system is still lacking. Overall, this study identifies the RsmY and RsmZ sRNAs as novel iron-responsive regulatory factors in *P. aeruginosa* static cultures. Taken together, our results demonstrate how static growth models allow for more thorough examination of virulence regulation in *P. aeruginosa*

## RESULTS

### Static growth promotes iron-regulated expression of the RsmY and RsmZ sRNAs in a PQS-dependent manner

To better understand the mechanism of iron-regulated expression of T6SS genes, we investigated the effects of iron on regulatory systems known to affect T6SS expression. We began by quantifying the impact of iron supplementation on levels of the RsmY and RsmZ sRNAs. Wild type *P. aeruginosa* strain PAO1 and the isogenic Δ*pqsA* mutant were grown in Chelex-treated and dialyzed tryptic soy broth (DTSB), an iron-limited medium used for previous iron regulatory studies in *P. aeruginosa* (23, 26), with or without supplementation of 100 µM FeCl_3_. Since iron regulation of T6SS genes was previously shown to occur only in static growth (23), we grew duplicate cultures with and without 250 rpm shaking. Expression of the RsmY and RsmZ sRNAs was significantly upregulated under iron-starved conditions in static but not shaking cultures (**Fig. 1**). Moreover, iron regulation in static growth was completely disrupted in the Δ*pqsA* strain that is defective in AQ production. Thus, iron supplementation represses expression of the RsmY and RsmZ sRNAs in an *pqsA*-dependent manner, and similar to what we previously observed with T6SS, this regulation is specific to static growth.

**Fig. 1:**
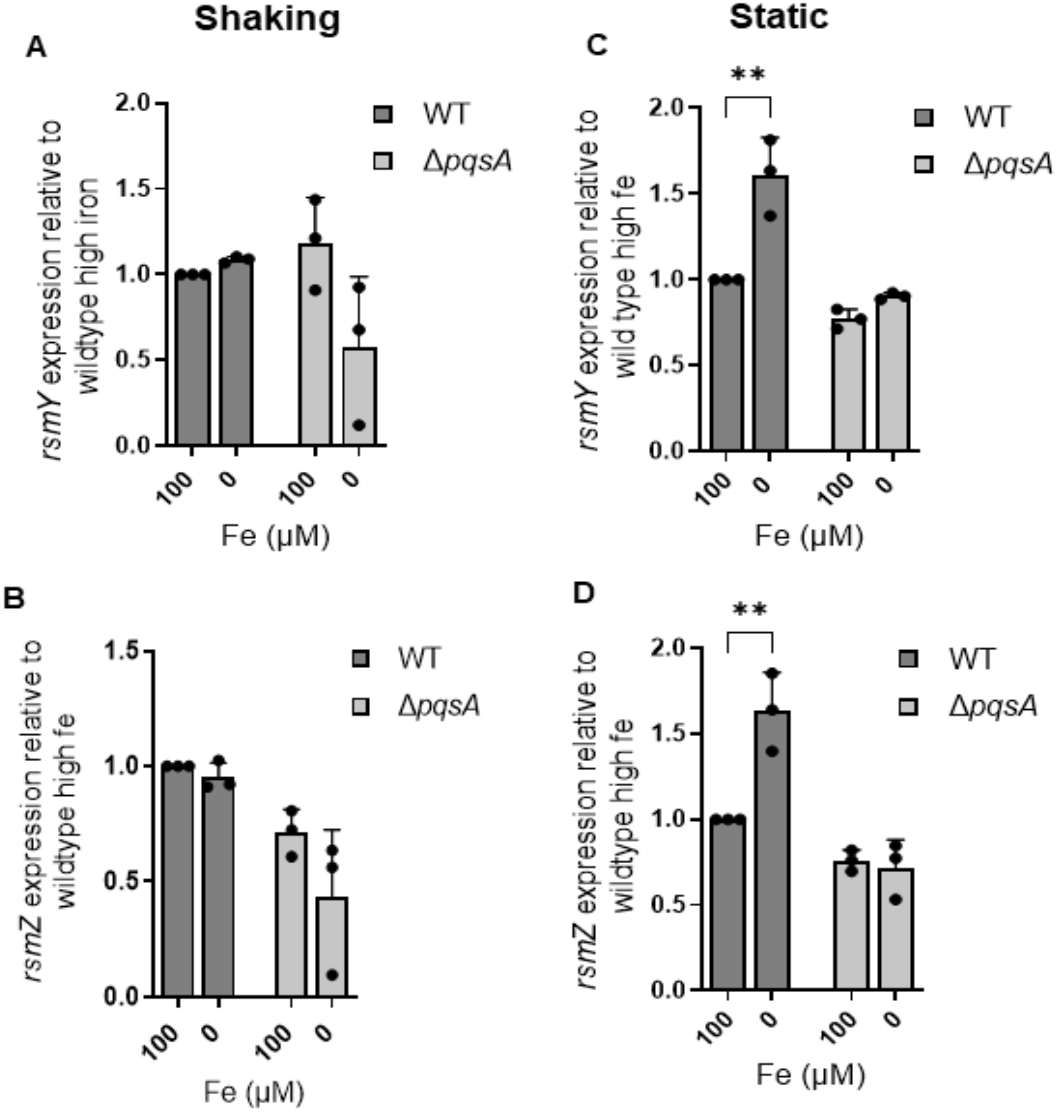
Static growth promotes expression of *rsmY* and *rsmZ* under iron-starved condition in a PQS-dependent manner: The indicated strains were cultured in DTSB and incubated for 18 h at 37°C supplemented with or without 100 μM of FeCl_3_ as the sole source of iron. The cultures were either incubated in shaking at 250 rpm (A and B) or static (C and D). RNA was isolated and reverse transcribed, and the resulting cDNA was analyzed by qPCR with primers and probes specific for RsmY and RsmZ. Results represent means ± SD of 3 independent experiments. Each biological replicate is indicated as data points shown in black solid circle. The asterisks indicate significant difference between indicated horizontal bars.** P <0.01, *** P <0.001 by Two-way ANOVA with Bonferroni’s post-test for multiple comparisons.

### Iron regulation of RsmY but not RsmZ is partially dependent on the *prrF* locus

Since the PrrF sRNAs are known to promote AQ production (26), we next determined if the Δ*prrF* mutant mediated iron-dependent regulation of RsmY and RsmZ. The wild type PAO1 and isogenic Δ*prrF* mutant were grown in DTSB with or without iron supplementation, in either shaking or static conditions, as above. Consistent with our previous data, iron starvation resulted in a significant increase in RsmY and RsmZ expression, and this induction was specific to static cultures (**Fig. 2**). Induction of RsmY was reduced but not eliminated in the Δ*prrF* mutant grown in static culture, while deletion of *prrF* had no significant impact on iron-regulated expression of RsmZ (**Fig. 2**). Thus iron-regulated expression of RsmY is partially dependent on the PrrF sRNAs, while RsmZ levels appear unaffected by PrrF regulation. The partial effect of PrrF on these sRNAs may be due to its indirect impact on AQ production, as *prrF* deletion reduces but does not eliminate production of AQ metabolites (23, 26).

**Fig. 2:**
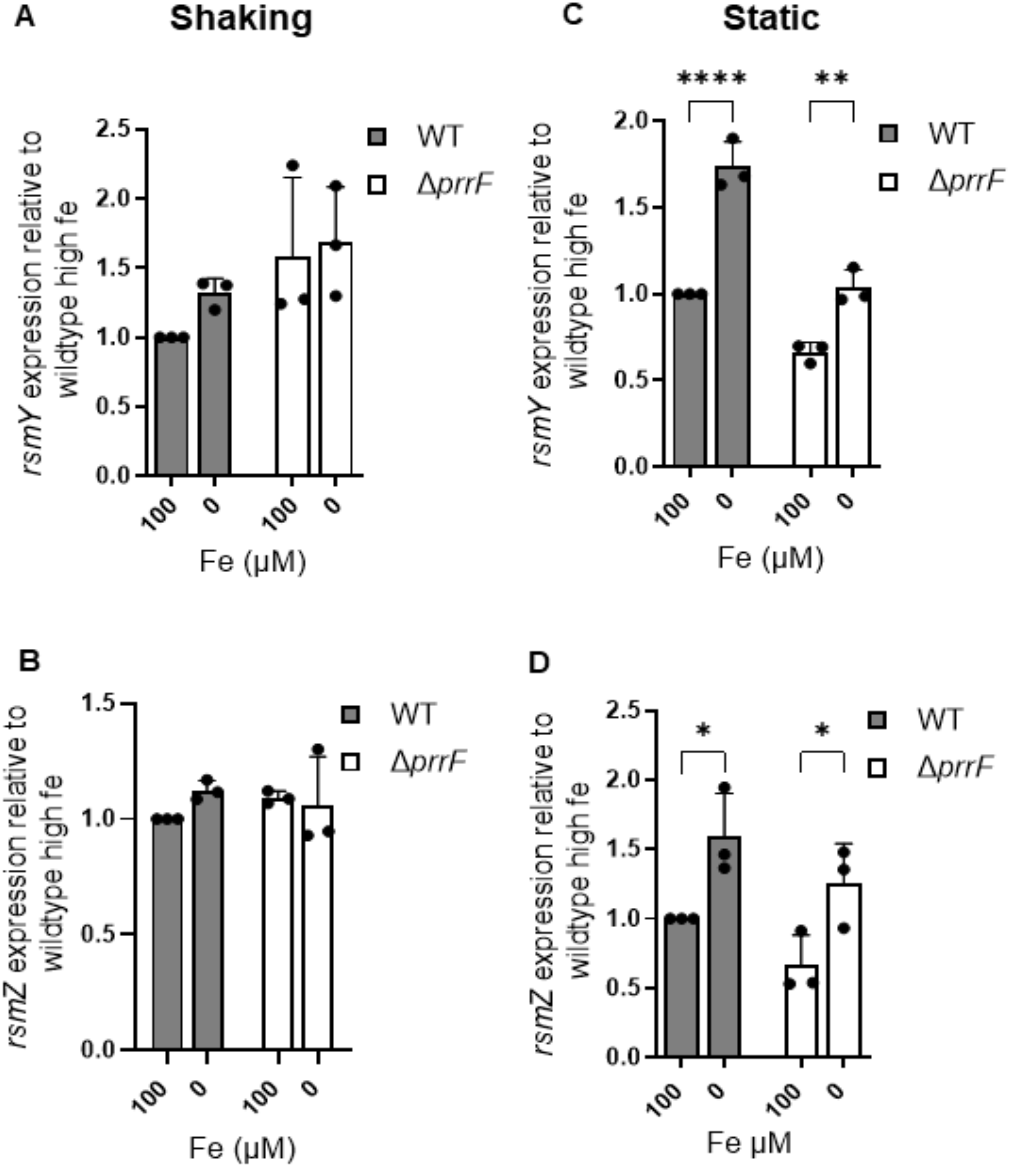
*PrrF* partially mediates upregulated expression of RsmY but not RsmZ in iron-starved static cultures: The indicated strains were cultured in DTSB and incubated for 18 h at 37°C supplemented with or without 100 μM of FeCl_3_ as the sole source of iron. The cultures were either incubated in shaking at 250 rpm (A and B) or static at 0 rpm (C and D). RNA was isolated and reverse transcribed, and the resulting cDNA was analyzed by qPCR with primers and probes specific for RsmY and RsmZ. Results represent means ± SD of 3 independent experiments. Each biological replicate is indicated as data points shown in black circle. The asterisks indicate significant difference between indicated horizontal bars.* P <0.05, ** P <0.01, *** P <0.001 by Two-way ANOVA with Bonferroni’s post-test for multiple comparisons.

### PqsA-dependent induction of RsmY and RsmZ by iron starvation occurs via the *rsmY* and *rsmZ* promoters

We next determined if iron starvation increased RsmY and RsmZ levels by increasing promoter activity of the *rsmY* and *rsmZ* genes. To test this, we fused each of the previously determined promoters (32, 33) with a promoterless *lacZY* reporter (**Fig. S1A-1B**) and introduced the resulting fusions into the chromosome of PAO1 and isogenic Δ*pqsA* mutant strains. In shaking cultures, iron starvation resulted in a slight decrease in β-galactosidase activity from the PAO1 *rsmY* reporter strain (**Fig. 3A**), while in static cultures *rsmY* promoter activity was induced nearly two-fold upon iron starvation (**Fig. 3C**). Similarly, static but not shaking culture allowed for increased activity of the *rsmZ* promoter under iron-depleted conditions (**Fig. 3B,D**). In agreement with the real time PCR results in **Figure 1**, iron regulation of both the *rsmY* and *rsmZ* promoters was ablated in the Δ*pqsA* mutant (**Fig. 3C-D**). Interestingly, we noted the activity of the *rsmZ* promoter was more strongly dependent on *pqsA* than that of the *rsmY* promoter in both shaking and static cultures, regardless of iron supplementation (**Fig. 3B,D**). Therefore, our data suggest that while iron-dependent regulation of both P_*rsmY*_ and P_*rsmZ*_ is AQ-dependent, the *rsmZ* promoter exhibits a stronger dependency on the AQ quorum sensing system.

**Fig. 3:**
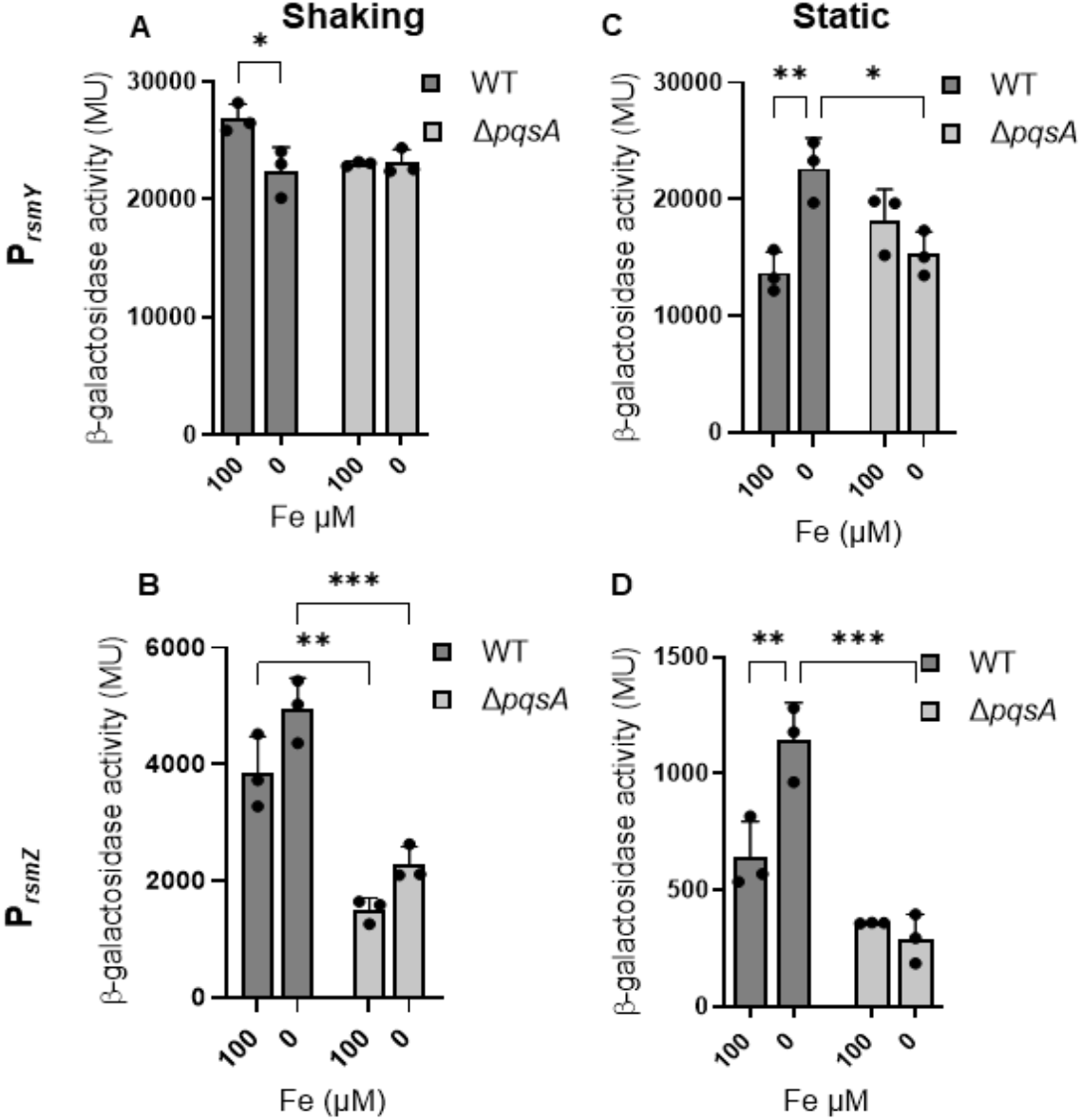
PqsA-mediated induction of RsmY and RsmZ occurs via the *rsmY* and *rsmZ* promoters. The indicated strains were cultured in DTSB and incubated for 18 h at 37°C supplemented with or without 100 μM of FeCl_3_ as the sole source of iron. The cultures were either incubated in shaking at 250 rpm (A and B) or static at 0 rpm (C and D). The indicated strains containing the *PrsmY*-lacZ or *PrsmZ*-lacZ transcriptional fusion were collected and subjected to β-galactosidase activity assay. Results represent mean ± SEM of 3 independent experiments. Individual data points are shown in black circle. The asterisks indicate significant difference between indicated horizontal bars.*P<0.05,** P <0.01, *** P <0.001 by Two-way ANOVA with Bonferroni’s post-test for multiple comparisons.

### PrrF is partially responsible for induction of *rsmY* and *rsmZ* promoter activity upon iron-starvation

We next tested the impact of *prrF* deletion on *rsmY* and *rsmZ* promoter activity by introducing the above reporters into the Δ*prrF* mutant. Consistent with reporter assays in **Figure 3**, iron starvation induced activity of both the *rsmY* and *rsmZ* promoters in the wild type strain in static but not shaking conditions (**Fig. 4**). In shaking conditions, deletion of the *prrF* locus had no significant impact on *rsmY* promoter activity (**Fig. 4A**), while activity of the *rsmZ* promoter was significantly reduced in the Δ*prrF* mutant grown in low iron shaking culture (**Fig. 4B**). This may reflect the stronger dependency of the *rsmZ* promoter on AQs (**Fig. 3**), production of which are dependent on the PrrF sRNAs in both shaking and static conditions (23). In contrast to the wild type grown in static conditions, static Δ*prrF* mutant cultures showed no significant iron-dependent regulation of *rsmY* or *rsmZ* promoter activity (**Fig. 4C,D**). Taken together, these data indicate that the PrrF sRNAs mediate, at least in part, iron-dependent regulation of the RsmY and RsmZ sRNAs under static conditions, likely through their effects on AQ production.

**Fig. 4:**
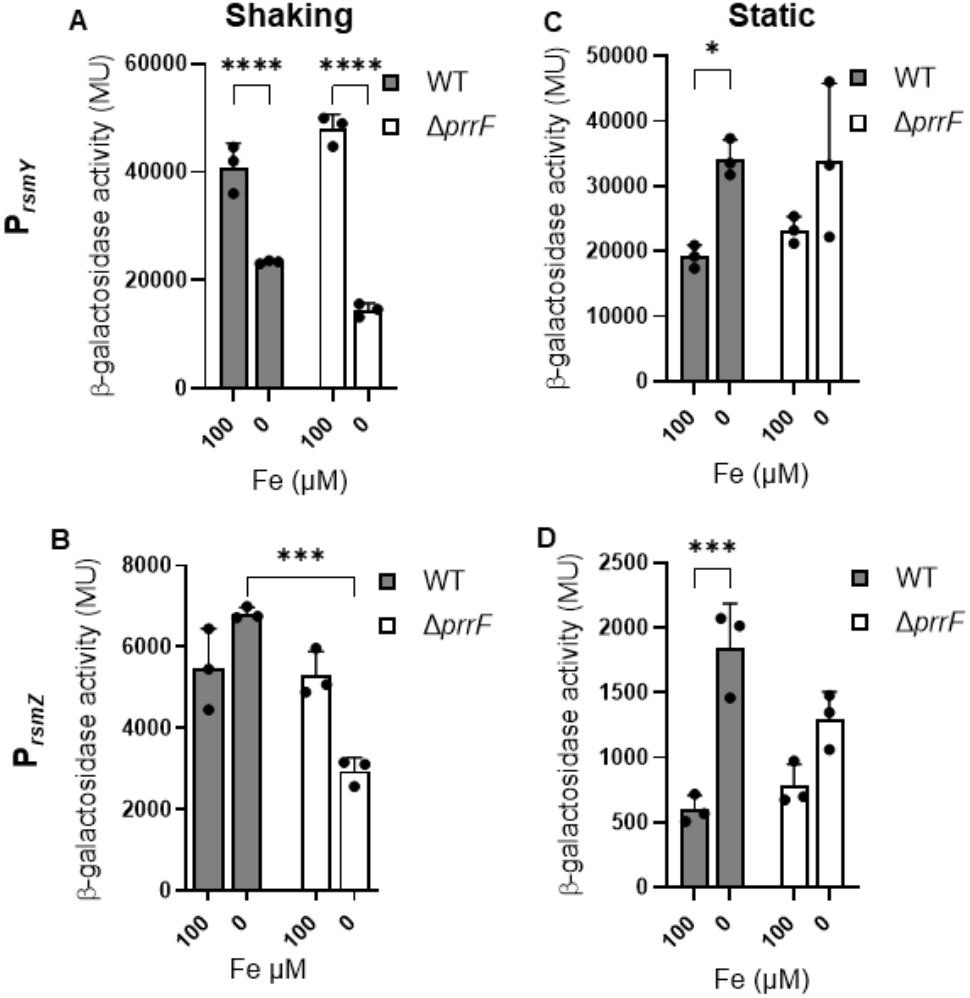
Differential iron-regulation *rsmY* and *rsmZ* under shaking and static cultures is partially dependent on the *prrF* locus: The indicated strains were cultured in DTSB and incubated for 18 h at 37°C supplemented with or without 100 μM of FeCl_3_ as the sole source of iron. The cultures were either incubated in shaking at 250 rpm (A and B) or static (C and D). The indicated strains containing the *PrsmY*-lacZ or *PrsmZ*-lacZ transcriptional fusion were collected and subjected to β-galactosidase activity assay. Results represent mean ± SEM of 3 independent experiments. Individual data points are shown in black circle. The asterisks indicate significant difference between indicated horizontal bars.*P<0.05,** P <0.01, *** P <0.001 by Two-way ANOVA with Bonferroni’s post-test for multiple comparisons.

### Iron regulation of RsmY is dependent on two putative PqsR binding sites in the *rsmY* promoter

Since our data above show PqsA-dependent iron regulation of the *rsmY* and *rsmZ* promoters, we identified palindromic sequences indicative of a binding site for the PqsR/MvfR transcription factor, the activity of which is dependent upon the PQS quorum sensing molecule (28). We found two such sites in the *rsmY* promoter, disrupted the palindrome by deleting sequences as shown in **Figure S1C**, and introduced the altered promoter fusions into wild type PAO1. As expected, we observed no significant induction of the wild type *rsmY* promoter from the wild type PAO1 strain grown in shaking conditions (**Fig. 5A**). Curiously, mutation of the *rsmY* promoter led to an increase in its activity in iron-replete, shaking conditions, and this increase was also observed with the wild type *rsmY* promoter in the Δ*pqsA* background (**Fig. 5A**). This increase was not observed in earlier experiments with the *pqsA* strain carrying the wild type *rsmY* reporter (**Fig. 3A**), thus it is possible this increase was due to experimental variation. In static growth conditions, iron limitation expectantly upregulated wild type *rsmY* promoter activity, while alteration of either or both of the predicted PqsR/MvfR binding sites eliminated iron regulation in static growth (**Fig. 5B**). This suggests that the palindromic sequences of P_*rsmY*_ are important for iron regulation of RsmY sRNA when *P. aeruginosa* is grown in static cultures.

**Fig. 5:**
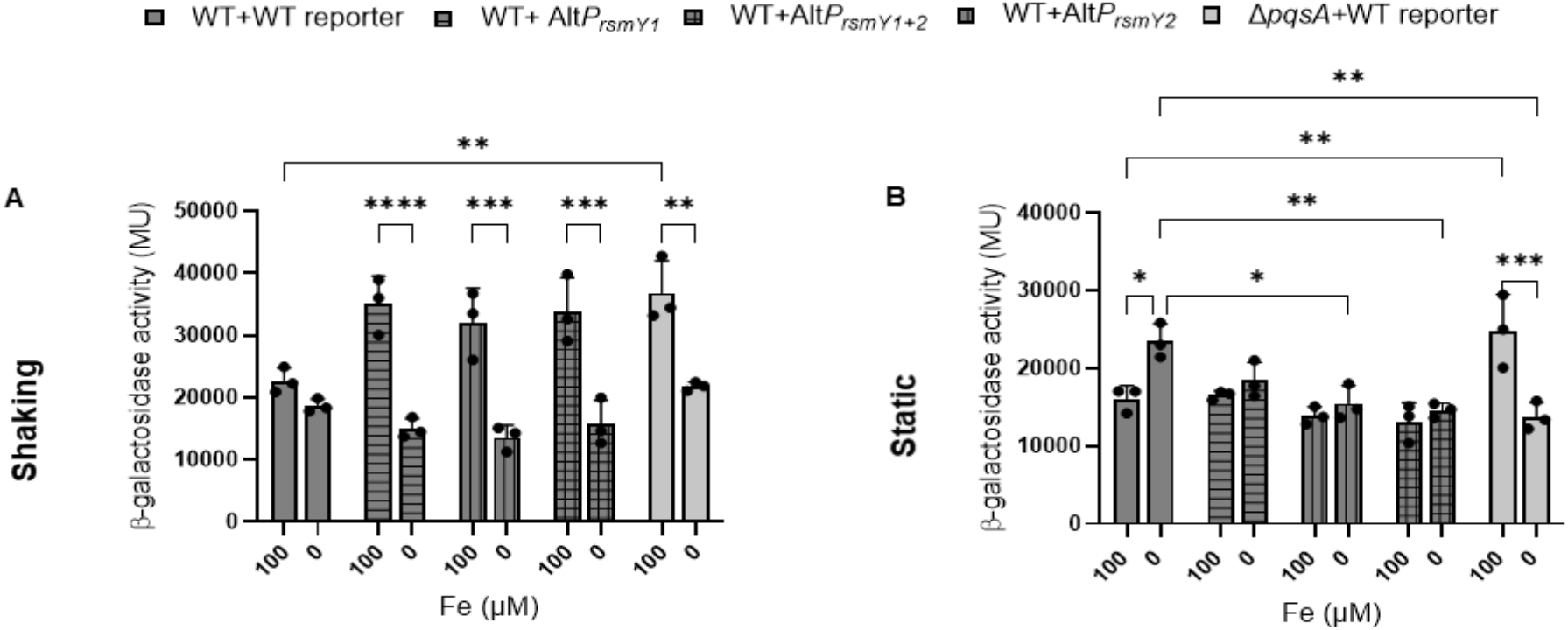
Iron-regulation of *rsmY* static cultures is dependent upon two palindromic promoter sequences. The indicated strains were cultured in DTSB and incubated for 18 h at 37°C supplemented with or without 100 μM of FeCl_3_ as the sole source of iron. The cultures were either incubated in shaking at 250 rpm(A) or static (B). The indicated strains containing the *P*_*rsmY1*_-lacZ or Alt*P*_*rsmY1*_-lacZ or Alt*P*_*rsmY2*_-lacZ or Alt*P*_*rsmY1+2*_-lacZ transcriptional fusion were collected and subjected to β-galactosidase activity assay. Results represent mean ± SEM of 3 independent experiments. Individual data points are shown in black circle. The asterisks indicate significant difference between indicated horizontal bars. *P<0.05,** P <0.01, *** P <0.001 by Two-way ANOVA with Bonferroni’s post-test for multiple comparisons.

### Altering a palindromic sequence of the *rsmZ* promoter disrupts PqsA-mediated iron-regulation only under static growth

We similarly altered a palindromic sequence in the promoter of *rsmZ* as shown in **Figure S1D**, and we moved this resulting reporter construct onto the chromosome of wild type PAO1. As previously seen, iron starvation had no significant induction effect on *rsmZ* promoter activity in either wild type or Δ*pqsA* when these strains were grown in shaking culture (**Fig. 6A**). Also as expected, iron starvation significantly induced *rsmZ* promoter from the wild type PAO1 strain when grown in static conditions, and this induction was dependent upon *pqsA* **(Fig. 6B)**. Notably, wild type PAO1 carrying the altered *rsmZ* showed drastically reduced promoter activity in both shaking and static conditions (**Fig. 6A-B**). This shows that the palindromic sequence in P_*rsmZ*_ has a dominant impact on its activity regardless of growth condition. Combined, these data suggest that PqsA-dependent iron regulation of *rsmZ* promoter activity requires the presence of a palindromic sequence in P_*rsmZ*_.

**Fig. 6.**
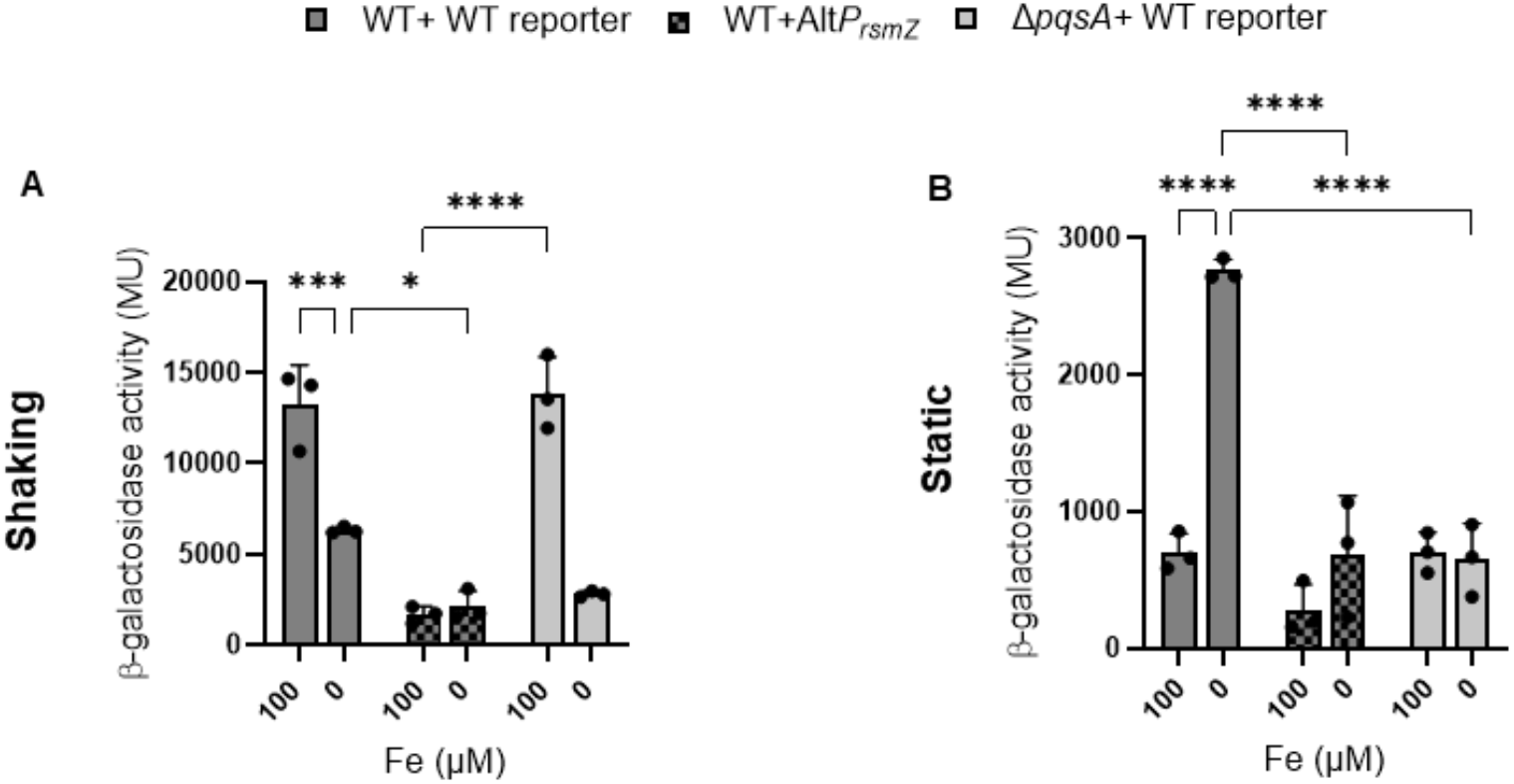
Iron regulation of *rsmZ* in static conditions occurs via a palindromic sequence in the promoter: The indicated strains were cultured in DTSB and incubated for 18 h at 37°C supplemented with or without 100 μM of FeCl_3_ as the sole source of iron. The cultures were either incubated in shaking at 250 rpm(A) or static at 0 rpm (B). The indicated strains containing the *P*_*rsmZ*_-lacZ or Alt*P*_*rsmZ*_-lacZ transcriptional fusion were collected and subjected to β-galactosidase activity assay. Results represent mean ± SEM of 3 independent experiments. Individual data points are shown in black circle. The asterisks indicate significant difference between indicated horizontal bars.*P<0.05,** P <0.01, *** P <0.001 by Two-way ANOVA with Bonferroni’s post-test for multiple comparisons.

### PqsA is required for iron-regulated promoter activity of the HSI-2 T6SS gene *hsiA2*

We next determined if iron-dependent regulation of T6SS gene expression likely occurred via upregulation of the RsmY and RsmZ sRNAs, which would sequester the RsmA RNA binding protein. Since this regulation would occur via a post-transcriptional regulatory event, we constructed transcriptional and translational fusions of the first gene in the HSI-2 T6SS locus, *hsiA2*. For the transcriptional fusion, we fused the known promoter of *hsiA2* to a promoterless *lacZY* operon as shown in **Figure S2A**. A combined transcriptional/translational reporter was also constructed by fusing the native *hsiA2* promoter and 5’ UTR to the promoterless *lacZY* operon (**Fig. S2B**). For the *hsiA2* translational fusion, we fused a constitutive *lac* promoter to the *hsiA2* 5’ UTR followed by the *lacZY* operon lacking its native RBS (**Fig. S2C**). We introduced each of the reporter constructs into the chromosomal *att* site of wild type PAO1 and the isogenic Δ*pqsA* mutant. Activity of the *hsiA2* transcriptional reporter from wild type PAO1 was induced 2.7 fold in static culture, as compared to only 1.4 fold induction in shaking condition (**Fig. 7A,D**). Significant iron regulation of the transcriptional fusion was eliminated in the Δ*pqsA* mutant when grown in static conditions (**Fig. 7D**), suggesting PqsA-dependent iron regulation of *hsiA2* under this condition occurs at the transcriptional level. The combined transcriptional-translational reporter showed similar results, with robust PqsA-dependent iron regulation of *hsiA2* occurring in static but not shaking conditions (**Fig. 7B,E**). Despite the known regulation of *hsiA2* by RsmA, however, we observed no significant iron regulation of the *hsiA2* translational reporter in the wild type strain grown in static conditions (**Fig. 7F**). Thus, our data suggest that iron regulation of *hsiA2* is mediated via PqsA via the *hsiA2* promoter.

**Fig. 7:**
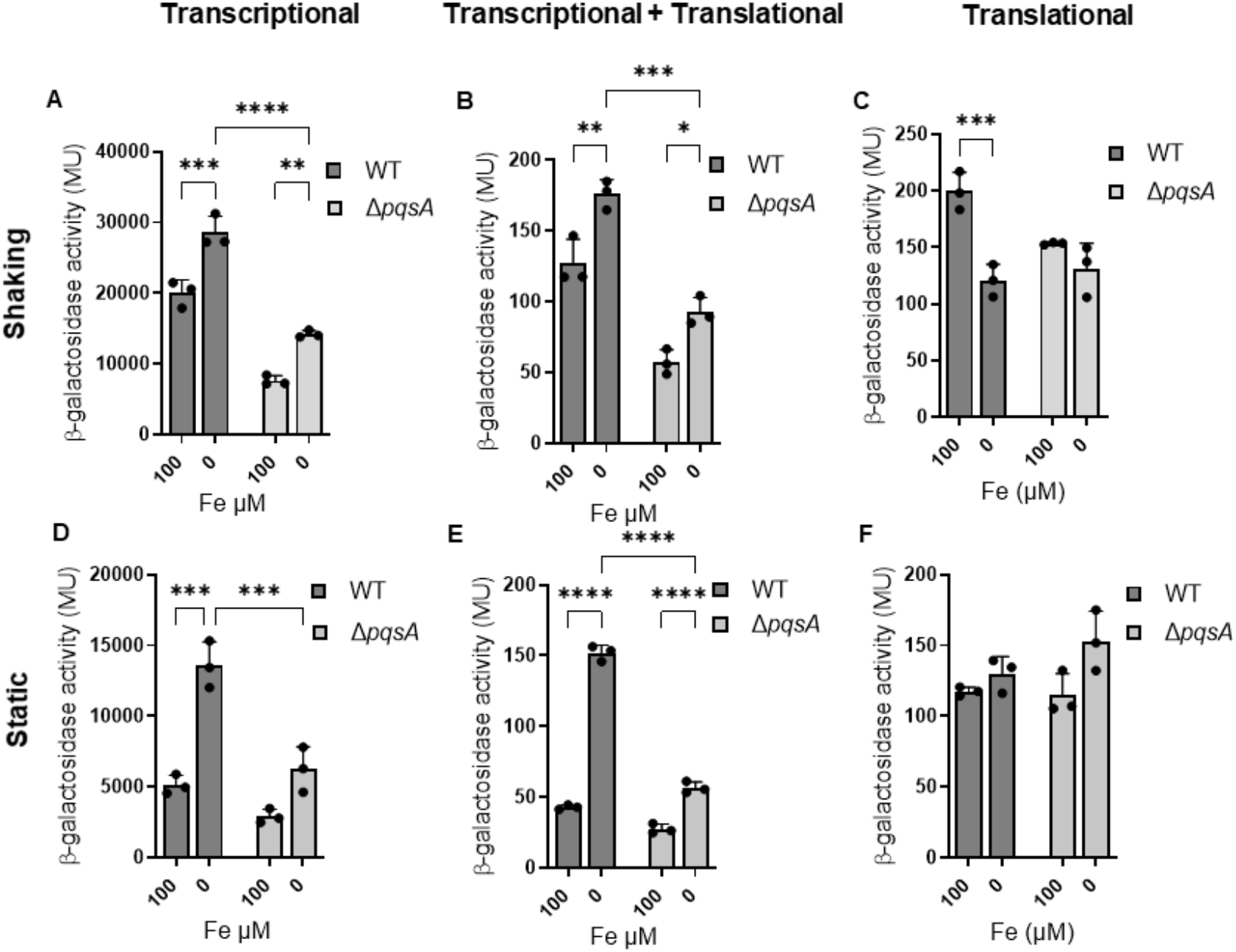
PqsA mediates promoter dependent iron regulation of T6SS gene *hsiA2*: The indicated strains were cultured in DTSB and incubated for 18 h at 37°C supplemented with or without 100 μM of FeCl_3_ as the sole source of iron. The cultures were either incubated in shaking at 250 rpm (A, B and C) or static (D, E and F). The indicated strains containing the *PhsiA2*-lacZ is transcriptional fusion (A and D), *PhsiA2+*UTR*hsiA2*-lacZ is transcriptional + translational fusion (B and E) and the translational fusions (C and F) consists of an in-built promoter *Plac*+UTR*hsiA2*-lacZ fused with untranslated region (UTR) of *hsiA2*. These fusions were introduced in WT PAO1 and Δ*pqsA*. They were harvested and subjected to β-galactosidase activity assay. Results represent mean ± SEM of 3 independent experiments. Individual data points are shown in black circle. The asterisks indicate significant difference between indicated horizontal bars.*P<0.05,** P <0.01, *** P <0.001 by Two-way ANOVA with Bonferroni’s post-test for multiple comparisons.

### PrrF is partially responsible for transcriptional iron regulation of *hsiA2*

We next examined the effect of PrrF on the activity of the transcriptional, translational, and combined reporters of *hsiA2* by moving the reporter constructs shown in **Figure S2** into the *att* site of the Δ*prrF* mutant. Static growth promoted consistent iron-dependent regulation of *hsiA2* promoter activity in wild type PAO1, and we observed a slight but significant reduction in activity of this reporter from the Δ*prrF* mutant when grown in static, low iron conditions (**Fig. 8A, D**). Curiously, the combined transcriptional-translational reporter showed significant induction by iron starvation in both static and shaking conditions, yet this effect was dependent on the *prrF* locus only in shaking conditions (**Fig. 8B, E**). We also observed a slight but significant upregulation of the *hsiA2* translational reporter activity in iron-depleted static culture, and this effect was dependent upon PrrF (**Fig. 8C**). Thus, our data suggest PrrF is at least partially responsible for iron-dependent regulation of the HS2-T6SS genes during static growth, though it remains unclear if this regulation is likely to occur through AQs, the Rsm sRNAs, or both.

**Fig. 8:**
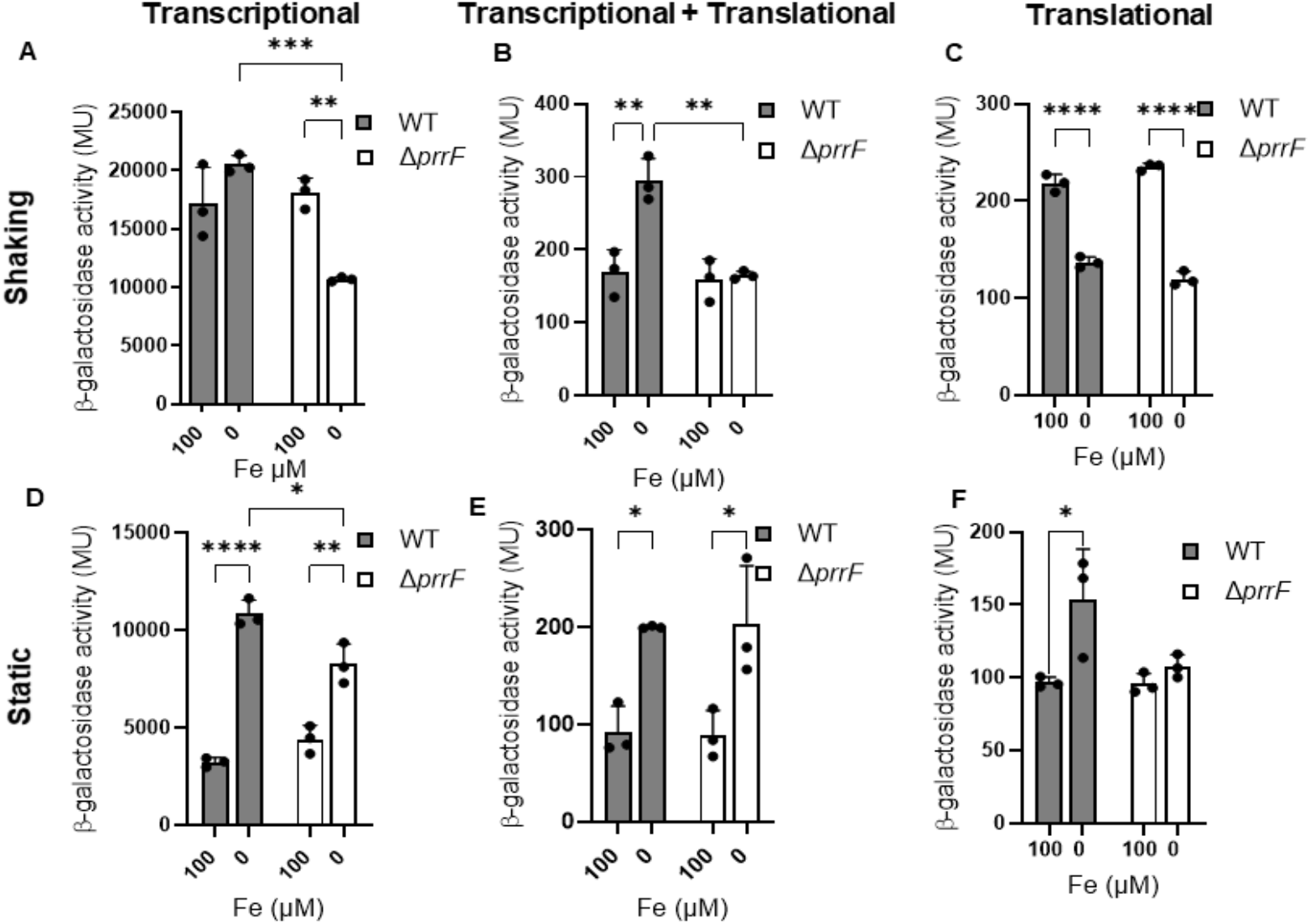
PrrF exerts transcriptional and posttranscriptional control on the expression of T6SS gene *hsiA2*: The indicated strains were cultured in DTSB and incubated for 18 h at 37°C supplemented with or without 100 μM of FeCl_3_ as the sole source of iron. The cultures were either incubated in shaking at 250 rpm (A, B and C) or static at 0 rpm (D, E and F). The indicated strains containing the *PhsiA2*-lacZ is transcriptional fusion (A and D), *PhsiA2+*UTR*hsiA2*-lacZ is transcriptional + translational fusion (B and E) and the translational fusions (C and F) consists of an in built promoter *Plac*+UTR*hsiA2*-lacZ fused with untranslated region (UTR) of *hsiA2*. These fusions were introduced in WT PAO1 and Δ*pqsA*. They were harvested and subjected to β-galactosidase activity assay. Results represent mean ± SEM of 3 independent experiments. Individual data points are shown in black circle. The asterisks indicate significant difference between indicated horizontal bars.*P<0.05,** P <0.01, *** P <0.001 by Two-way ANOVA with Bonferroni’s post-test for multiple comparisons.

## Discussion

The chronically infected CF lung is a constantly changing environment, and changes in iron availability are known to correlate with CF lung disease progression from acute to chronic stages of *P. aeruginosa* infection (34-36). *P. aeruginosa* has a strong metabolic requirement for iron and as such, it must rapidly adapt to changing iron availability in the lung. Iron starvation has long been known to regulate virulence gene expression in *P. aeruginosa*, though historically these studies have been performed in homogenous, shaking cultures, which are unlikely to represent growth conditions in the CF lung. Static growth may provide a more biologically relevant in vitro model for understanding gene regulation in chronically infected CF lungs. Supporting this idea, we previously showed that static growth allowed for iron regulation of several systems not previously shown to be iron-responsive (23), including the HSI-2 T6SS that is closely associated with chronic infection (22, 37). In the current study, we show that iron starvation induces expression of the RsmY and RsmZ sRNAs, which have previously been shown to promote expression of genes for the HSI-2 T6SS (20). We further show that this regulation is specific to static growth, is completely dependent on PqsA, and is partially dependent upon the PrrF sRNAs, also similar to what was previously observed for T6SS (23). Moreover, we found that PqsA-dependent iron regulation of *hsiA2*, the first gene in the HSI-2 T6SS gene cluster, occurs primarily at the promoter level, while PrrF-dependent effects were observed for both the transcriptional and translational *hsiA2* reporters. This last finding was surprising, as we had hypothesized that the Rsm sRNAs post-transcriptionally mediated iron-dependent regulation of T6SS genes. Nevertheless, these findings point to a novel role for the Rsm sRNAs in *P. aeruginosa’s* response to iron starvation during growth in static conditions.

Since iron regulation of the RsmY and RsmZ sRNAs was dependent on PqsA, we identified potential PqsR/MvfR binding sites in the *rsmY* and *rsmZ* promoters. Identification of these sites was based on previous literature showing that PqsR/MvfR binds to a palindromic sequence known as a LysR box (28, 38). Disruption of the palindromic sequence in P_*rsmZ*_ affected its expression regardless of whether the strains were grown in shaking or static conditions (**Fig. 6**). Moreover, activity of the altered *rsmZ* promoter was similar to that of the wild type promoter in the Δ*pqsA* background when grown in static conditions **(Fig. 6)**. Combined, these data suggest that PqsA-dependent iron regulation of RsmZ is dependent on the PqsR/MvfR protein, in the presence of AQs, binding to this palindromic sequence. In contrast, altering the palindromic sequences of the *rsmY* promoter did not mimic activity of the wild type *rsmY* promoter in the Δ*pqsA* strain when grown in static conditions (**Fig. 5**), suggesting PqsA-dependent iron regulation of RsmY is more complex than that of RsmZ. Combined, our data show that iron starvation results in PqsA-dependent transcriptional control of the RsmY and RsmZ sRNAs, yet the mechanism for how each of these sRNAs is affected by PqsA appears to be distinct.

Since *ΔpqsA* lacks AQ production, we presume that expression of the RsmY and RsmZ sRNA levels are AQ-dependent. However, there are multiple classes of AQs, including the AHQ and PQS metabolites that can both function as quorum sensing molecules (28). Previous work in our lab shows that the levels of some AHQs like 4-hydroxy-2-heptylquinoline (HHQ) and 2-nonyl-4-hydroxy-quinoline (NHQ) are much higher in static versus shaking cultures, while the C7 and C9 congeners of 3,4-dihydroxy-2heptylquinoline (PQS) are reduced in static cultures (23). HHQ and PQS can both promote PqsR/MvfR interaction with targeted promoters (28). However, AHQs and PQS have been shown to exert differential effects on PqsR/MvfR-dependent gene expression (28, 31), suggesting that different AQs may impact PqsR/MvfR affinity or interaction with different promoter sequences. In other words, differences in PqsA-dependent iron regulation of RsmY and RsmZ in static versus shaking conditions could be due to varying levels of distinct AQ species in each condition. Cell-to-cell transmission of these different AQ classes also likely varies, as PQS promotes the production of extracellular vesicles that can also deliver AQs (39). In this vein, it is possible that the differential impact of PqsA on the expression of Rsm sRNAs in shaking and static cultures could be due to the perturbation in shaking cultures that affects physiologically-relevant delivery of AQs via extracellular vesicles.

As we know that AQ production is indirectly promoted by the iron-responsive PrrF sRNAs, we speculated that the PqsA-dependent iron-regulation of Rsm would be dependent on the *prrF* locus. In agreement with this idea, we found that iron-regulated expression RsmY levels and *rsmY* promoter activity was partially disrupted in the Δ*prrF* strain (**Fig. 2, 4**). Similarly, iron-regulated *rsmZ* promoter activity was modestly affected by *prrF* deletion, though RsmZ RNA levels were not significantly impacted **(Fig. 2, 4)**. The partial effect elicited by Δ*prrF* deletion is likely due to its indirect effect on AQ production, which results in decrease levels of AQs, while the Δ*pqsA* mutant produces no detectable AQs (40). Thus our model currently posits that iron regulation of RsmY and RsmZ occurs via PrrF’s positive impact on production of AQs, which in turn activate expression of the RsmY and RsmZ sRNAs.

Since T6SS is regulated by the Rsm system in a post-transcriptional manner (12, 20), we used transcriptional and translational reporter fusions of *hsiA2* to examine the level at which iron affects its expression. We demonstrated that PqsA and PrrF promote iron-regulated *hsiA2* transcriptional reporter activity under static conditions. Surprisingly, we did not observe any significant changes in the activity of the *hsiA2* translational reporter, despite the known regulation of *hsiA2* by RsmA. This could suggest that PqsR/MvfR is capable of directly interacting with the *hsiA2* promoter, and that iron-regulated expression of RsmY and RsmZ does not play a significant role in iron regulated expression of HSI-2 T6SS. We also observed that PrrF mediated a slight but significant induction of translational activity of *hsiA2*, suggesting partial PrrF dependent control this gene in static conditions. This data aligns with our proteomics data where we established PrrF dependent iron repression of HSI-2 T6SS genes (23). Notably, our studies on T6SS gene expression were focused on the *hsiA2* promoter and UTR, and many questions remain regarding how individual genes in the T6SS are regulated at the transcriptional, translational, and post-translation level to affect T6SS function. Thus further studies are needed dissect the mechanisms by which iron, PrrF, and PqsA affect production of the HSI-2 T6SS.

While the current study was started in an effort to trace the mechanism behind iron regulated T6SS gene expression in static cultures, we for the first-time have identified iron regulation of the Rsm system in *P. aeruginosa*. Furthermore our studies implicated the PQS quorum sensing system in the iron regulation of the Rsm system. Static cultures may better reflect conditions within the host, as bacterial communities are able to establish biochemical gradients that affect their metabolism, nutrient acquisition, and ability to coexist. Moreover, static growth is likely to promote contact-dependent interactions, which are required for T6SS function, as well as surface attachment and initiation of biofilm development. As such, our study provides insight into the concept of using biologically more relevant condition to study the impacts of iron regulation, and has yielded a novel iron-responsive regulatory in this important human pathogen. Further investigations are required to dissect the complex environmental factors that affect iron regulation in static bacterial communities, and we expect this work will provide further insight into this complex yet exciting network of physiological processes. This idea could be utilized to generate knowledge that has a broader spectrum in understanding bacterial regulation and finding new therapeutic targets that are relevant during chronic infection.

## MATERIALS AND METHODS

### Bacterial strains and growth conditions

Bacterial strains used in this study are listed in **Table S1 (Supplementary Materials)**. *P. aeruginosa* lab strain PAO1 was routinely grown overnight by streaking from freezer stocks in tryptic soy agar (TSA) (Sigma, St Louis, MO) plates having 15g/L agar. Five isolated colonies were taken from each streaked plate and inoculated in 1.5 ml of TSB. For iron starvation experiments, bacterial strains were sub-cultured into Chelex-treated and dialyzed trypticase soy broth (DTSB) prepared as previously described(23). This media was either supplemented with 0 μM or 100 μM FeCl_3._ The cultures were incubated at a shaking rate of either 250rpm (shaking conditions) or 0rpm (static conditions) for 18 hours at 37°C. *E. coli* strains were grown in LB (Sigma, St Louis, MO) and streaked on LB plates having 15g/L agar when applicable. The antibiotics were added in the following concentration ampicillin, 100 μg/ml; tetracycline, 10 or 15 μg/ml, gentamycin 20 μg/ml; for *E. coli* and carbenicillin, 250 μg/ml; tetracycline, 150 μg/ml, and gentamycin 50 μg/ml for PAO1.

### Construction of reporter strains

The promoter region of *rsmY* and *rsmZ* sRNA were amplified by PCR using primers mentioned in **Supplementary Table S1** and the PAO1 genomic DNA as a template. The PCR product was cloned in a TA cloning vector PCR2.1 (Invitrogen) and confirmed by sequencing. The promoter fragment was then digested by restriction enzymes EcoR1 and BamH1 to get them cloned into a promoter-less *lacZY* reporter plasmid Mini-CTX-*LacZY* to generate Mini-CTX-P_*rsmY*_-*LacZY* and Mini-CTX-P_*rsmZ*_-*LacZY* plasmid for transcriptional reporter assays. The vectors were digested by the same restriction enzymes used for the promoter fragment. Similarly, we generated transcriptional reporter plasmid for HSI-2 core gene cluster mRNA of T6SS by PCR amplifying the promoter region upstream of *hsiA2*, the first gene of the cluster and cloning it into Mini-CTX-*LacZY* to generate Mini-CTX-P_*hsiA2*_-*LacZY* plasmid for transcriptional reporter assay of T6SS core cluster of HSI-2 mRNA. We further generated transcriptional + translational **(Fig.S2B)** reporter of *hsiA2* by using Mini-CTX-*lacZY*-SD vector, which lacks the *lacZ* Shine-Dalgarno site, was constructed as mentioned in our previous work(26). The leader and the promoter sequence of *hsiA2* that extends from 150 nt upstream of +1 transcriptional start site to 15 nt downstream of the translational start site was PCR amplified using oligonucleotides mentioned in the **Supplementary Table S1**. The PCR product was cloned in a TA cloning vector PCR2.1 and the fragments were then digested by restriction enzymes EcoR1 and BamH1 and ligated with Mini-CTX-*lacZY*^-SD^ vector to generate Mini-CTX-P_*hsiA2*_ *-lacZY*^-SD^ plasmid. Similarly, we also constructed translational reporter by by PCR amplification of the *hsiA2* 5’ UTR, from the +1 transcriptional site to 15 nt downstream of the translational start site, using oligonucleotides mentioned in the **Supplementary Table S1** in which the PCR product was similarly cloned into Mini-CTX-P_*lac*_-*lacZY*^-SD^ that contains a heterologous promoter (P_*lac*_) to generate Mini-CTX-P_*lac*_-UTR_*hsiA2*_-*lacZY*^-SD^. Altered *rsmY and rsmZ* reporters were constructed using the QuikChange II XL site-directed mutagenesis kit (Agilent) following the manufacturer’s instructions, with PCR2.1-*rsmY* and PCR2.1-*rsmZ* as the template and primers mentioned in **Supplementary Table S1**. All the reporter constructs were introduced into PAO1, Δ*pqsA* and Δ*prrF1,2* chromosomes by integrating them at the *att* site as previously described(41).

### Real time PCR analysis

Quantitative real time PCR (qRT-PCR) analysis was of each of the strains of wild type *Pseudomonas aeruginosa* PAO1 and the isogenic deletion mutants Δ*prrF*, or Δ*pqsA* grown in DTSB supplemented with (high iron) or without (low iron) 100 µM FeCl_3._ The cultures were incubated at a shaking rate of either 250rpm (shaking conditions) or 0rpm (static conditions) for 18 hours at 37°C. RNA extraction was done as mentioned in (luke ref). Briefly, *P. aeruginosa* cells were lysed after 18 hours using 2.5mg/mL lysozyme and incubated at 37°C for 30 minutes and RNA was extracted using the RNeasy mini kit (QIAGEN, Germantown, MD). DNase treatment was done twice for better DNA removal using Qiagen RNase-free DNase during RNA extraction and using NEB DNase after RNA extraction. cDNA synthesis was done with reverse transcription kit (Promega) using 50ng of total RNA. Real-time qualitative PCR analysis was performed as described previously(42) using an Applied Sciences Step One Plus Real Time PCR System (Life Technologies, Carlsbad, CA).Primers and probes were designed using IDT and are mentioned in supplementary material (**Table:S1**). Quantitation of cDNA was carried out using standard curves generated by performing qPCR of individual gene targets using cDNA reverse-transcribed from serial dilutions of RNA. *P. aeruginosa* expression data was normalized to *oprF* expression in aerobic shaking conditions, or *omlA* in static growth condition, as expression of these genes is not impacted by iron in each respective growth condition (42).

### β-Galactosidase activity

β-Galactosidase activity was measured as described previously(26). Briefly, bacteria were collected to determine the absorbances (A_600_), and cells were harvested by centrifugation and resuspended in Z buffer, followed by 1:10 dilution in Z buffer. Z buffer were prepared as previously described^26^. The cells were lysed using chloroform and 0.1% sodium dodecyl sulfate (SDS). ONPG (o-nitrophenyl-β-D-galactopyranoside) substrate was added to the solution, and the reaction was stopped with sodium carbonate after 10 to 40min, until a clear color change was observed. The reaction mixtures were briefly centrifuged, and the absorbance (A420) of the supernatant was determined. The β-galactosidase activity was calculated in Miller units: (1,000 X A420)/ (time [minutes] X volume [milliliters] X A600).

### Statistical analysis

Statistically significant changes were identified in all the experiments using two-way ANOVA on Prism 9, using Bonferonni’s posttest for multiple comparisons, with a significance threshold of *p* value < 0.05. Each of the experiments were conducted on at least three biological replicates and the results were represented as mean ± SD.

